# The epigenetic landscape in purified myonuclei from fast and slow muscles

**DOI:** 10.1101/2021.02.04.429545

**Authors:** Mads Bengtsen, Ivan Myhre Winje, Einar Eftestøl, Johannes Landskron, Chengyi Sun, Diana Domanska, Douglas P. Millay, Leonardo Meza-Zepeda, Kristian Gundersen

## Abstract

Muscle cells have different phenotypes adapted to different usage and can be grossly divided into fast/glycolytic and slow/oxidative types. While most muscles contain a mixture of such fiber types, we aimed at providing a genome-wide analysis of chromatin environment by ChIP-Seq in two muscle extremes, the almost completely fast/glycolytic extensor digitorum longus (EDL) and slow/oxidative soleus muscles. Muscle is a heterogeneous tissue where less than 60% of the nuclei are inside muscle fibers. Since cellular homogeneity is critical in epigenome-wide association studies we devised a new method for purifying skeletal muscle nuclei from whole tissue based on the nuclear envelope protein Pericentriolar material 1 (PCM1) being a specific marker for myonuclei. Using antibody labeling and a magnetic-assisted sorting approach we were able to sort out myonuclei with 95% purity. The sorting eliminated influence from other cell types in the tissue and improved the myo-specific signal. A genome-wide comparison of the epigenetic landscape in EDL and soleus reflected the functional properties of the two muscles each with a distinct regulatory program involving distal enhancers, including a glycolytic super-enhancer in the EDL. The two muscles are also regulated by different sets of transcription factors; e.g. in soleus binding sites for MEF2C, NFATC2 and PPARA were enriched, while in EDL MYOD1 and SOX1 binding sites were found to be overrepresented. In addition, novel factors for muscle regulation such as MAF, ZFX and ZBTB14 were identified.

## Introduction

The phenotype of skeletal muscle fibers differs as an adaption to different tasks. Some fibers have short twitches and rapid shortening velocities, but low endurance. Such fibers are used for short, but explosive external work (sprinting, throwing, stumbling etc.), but are easily fatigued and not very energy efficient. Other fibers have slow twitches and shortening velocities, but are fatigue resistant and have a low energy expenditure. Such fibers are used for tasks as keeping body and limb posture.

Fibers are generally classified into “fiber types” related to the predominant myosin heavy chain (MyHC) isoenzyme expressed in the cell. MyHC is the strongest molecular determinant of shortening speed. In rodents there are four different MyHC genes expressed in adult limb muscles namely *Myosin Heavy Chain 7* (*Myh7*), *Myh2, Myh1* and *Myh4* giving rise to the fiber types 1 (slowest), 2A, 2X and 2B (fastest) respectively. Partly coupled to the MyHC fiber type, the fibers display a spectrum of metabolic properties, from highly oxidative mitochondria-rich cells (type 1) to cells that are mainly glycolytic (type 2B). At the molecular level the different fiber types vary in isoform expression of various of proteins such as calcium pumps, oxygen related proteins and also sarcomere components other than MyHC.

Skeletal muscle is the most important metabolic organ in the body. It has been known for 40 years that differences in muscle fiber type composition is a strong individual predictor for developing metabolic syndrome (1). Metabolic syndrome is a cluster of conditions increasing the risk of heart disease, stroke and type 2 diabetes. These conditions include increased blood pressure, high blood sugar levels, excess body fat around the waist, and abnormal cholesterol or triglyceride levels. Metabolic syndrome is on the rise, and in several countries the prevalence is now over ¼ of the population (2). Epigenetic mechanisms like histone modifications and chromatin structure have been suggested to play an important role in the development of and predisposition for metabolic syndrome (3), but data supporting this is currently scarce.

Muscle fibers are post-mitotic and represent an interesting example of a balance between long term phenotypic stability yet malleability. Thus, phenotype can be changed by altering the pattern of electrical activity elicited in their sarcolemma by the motor neurons or electrical stimulation (4) and also by the mechanical stress created by contraction (5). These two external factors are the major mechanistic foundations for the effects of exercise on muscle. However, adult phenotype also dependents on the embryonal cell lineage and this origin limits the adaptive ranges of training effects (6). The transition process has been studied in detail in rats stimulated with electrical patterns mimicking extreme, but well-defined training (4). It appears that some traits require very long-term treatment in order to be altered e.g. type 2 to type 1 transformations, which might take more than 3 months (7), in contrast changes within type 2, i.e. 2B>2A>2X (8) and changes in metabolic and calcium sequestering enzymes can be altered more rapidly (9). Suggesting that epigenetic mechanisms may play an important role in the regulation of muscle plasticity (10, 11).

Recently the existence of a long term cellular muscle memory was demonstrated (12, 13), and in addition to a permanently elevated number of myonuclei (14, 15) epigenetic mechanisms might be involved (16). Epigenetic mechanisms might also be related to the observation that some individuals seem to have less malleable muscles than others, i.e. exercise resistance (17-19).

Since the overall chromatin environment and the modifications of the histones represent a form of overreaching gene regulation mechanism with the potential of long-lasting stability (16, 20, 21), we set out to compare chromatin environment in an extremely fast/glycolytic muscle (the m. extensor digitorum longus, EDL) and an extremely slow/oxidative muscle (the m. soleus) in mice.

The majority of studies on epigenetics are on tissue culture cells, and less has been done on tissue homogenates where results seem harder to interpret. In muscle, electron microscopy (22) and a specific marker for myonuclei (22-24) has revealed that only approximately 40 – 50% of the nuclei found in muscle tissue are myonuclei and since accounting for cellular heterogeneity is critical in epigenome-wide association studies (25-28) we aimed at purifying the myonuclei proper.

We recently reported that the nuclear envelope protein Pericentriolar material 1 (PCM1) can be used as a specific light microscopy marker to discern the skeletal muscle myonuclei in both rodents and humans (24), and we show here that this marker can be used to sort myonuclei to >95% purity before a subsequent epigenetic analysis. Furthermore, we show that the purification is necessary for the epigenetic landscape faithfully to reflect known features of muscle function, and the results indicate that purification should be used in studies aimed at elucidating the role of epigenetic mechanisms in muscle differentiation and plasticity.

## Results

### Purification of myonuclei allows a genome-wide analysis of muscle fiber specific chromatin

#### PCM1 labelling is specific for myonuclei isolated from the muscle tissue

To remove interference from non-myofiber nuclei in the tissue, we took advantage of an antibody against Pericentriolar material 1 (PCM1) which we have previously reported to selectively label myonuclei in skeletal muscle fibers and not satellite cells or stroma cells (24, 29) (Fig. S1A).

To further prove the specificity of PCM1 for muscle myonuclei we used a transgenic mouse model expressing Histone-2B coupled to GFP (H2B-GFP) under the control of the *skeletal muscle Actin Alpha 1* (*ACTA1*) promoter giving specific expression of the fusion construct in skeletal muscle fiber nuclei across fiber types (30). Single fiber analysis showed that H2B-GFP and PCM1 labelling invariably colocalized (Fig 1A-B and Fig. S1C). In some cases, DAPI labelled nuclei that were negative for both H2B-GFP and PCM1 were observed (arrows in Fig. 1D-E), probably representing non-myonuclei sticking to the fibers after isolation.

**Fig. 1:**
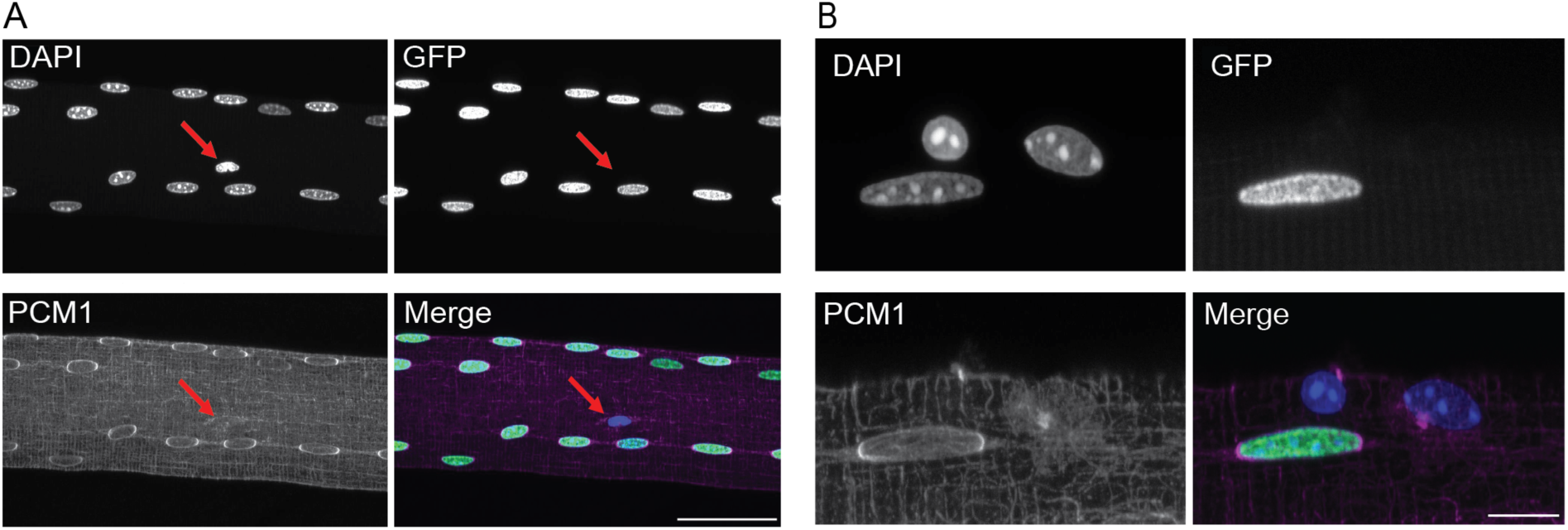
PCM1 selectively labels myonuclei in isolated single fibers. A) Max intensity Z projection of single fiber from transgenic mouse expressing the H2B-GFP (Green) construct under the control of ACTA1 stained against PCM1 (magenta). Counterstained with DAPI to visualize DNA (Blue) D) Arrow: nucleus negative for both PCM1 and H2B-GFP. Scale bar 50 *μ*m E) High resolution image of single fiber showing GFP positive and negative nuclei. Scale bar 10 *μ*m.

#### PCM1 can be used to isolate myonuclei for subsequent analysis across species

Flow cytometry analysis of nuclei from skeletal muscle displayed a 98.0 % overlap between GFP and PCM1 labelling of the nuclei in the transgenic mouse line (Fig. 2A-F).

**Fig. 2:**
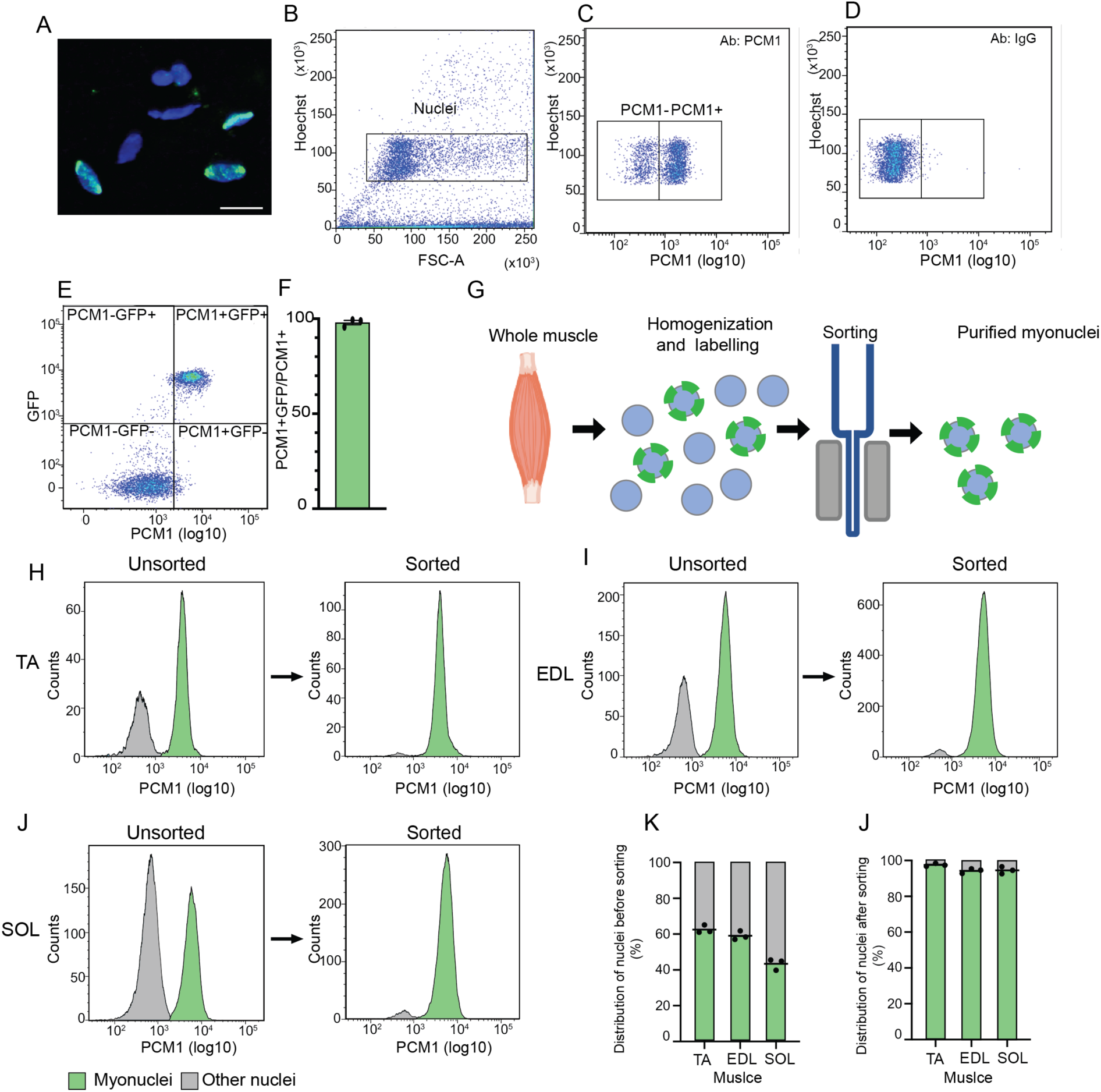
PCM1 can be used to isolate the myonuclei population from whole skeletal muscle tissue. A) Nuclei from whole muscle lysate stained with PCM1 antibody (green) and Hoechst to visualize DNA (blue), Scale bar 20 *μ*m. B) Representative scatterplot showing identification of single nuclei from skeletal muscle tissue by flow cytometry. C-D) Representative scatterplot of nuclei labeled with PCM1 and IgG, respectively. E) Representative scatterplot of co-localization between nuclei expressing H2B-GFP and positive for PCM1 in tibialis anterior (TA) from mouse expressing the H2B-GFP construct under the control of ACTA1, analyzed by flow cytometry. F) Co-localization between PCM1 and GFP positive nuclei in TA in the transgenic mouse model (n=3), error bare SEM. G) Workflow used to isolate the myo-specific nuclei from skeletal muscle. H-J) Representative histograms of nuclear distribution and magnetic sorting efficiency for the three muscles TA, EDL and soleus (SOL) from mice by flow cytometry. K) Quantification of nuclear distribution in full tissue (n=3). J) Quantification of sorting efficiency (n=3).

Analysis by flow cytometry of the nuclear composition in the tibialis anterior (TA) with mixed fiber type composition and the two muscle extremes EDL and soleus with a fast and slow fiber type composition, showed that the number of myonuclei varied from 40 % -60 % with the lowest number of myonuclei in the slow soleus (Fig. 2G-K), confirming that without sorting roughly half of the nuclei isolated from muscle homogenates are from other cell types than muscle fibers and might confound a genome-wide analysis aimed at this cell type.

To isolate the myonuclear fraction from the nuclei originating from the other cell types in the tissue we used a magnetic-assisted sorting approach (Fig. 2G-J). Using this method, we were able to isolate the myonuclei with a very high purity on an average of 95 % for all the three different muscle type (Fig. 2G-K). The same high sorting purity was obtained when the three same muscle types from rat were analyzed (Fig. S2).

#### Purification of nuclei is required for a valid genome-wide analysis of the myo-specific epigenome

To investigate the importance of nuclear purification for epigenetic analysis of muscle cells we performed chromatin immunoprecipitation on the PCM1 purified nuclei followed by next-generation sequencing (ChIP-Seq) in soleus and EDL using an antibody against Histone H3 acetylated at Lysine 27 (H3K27ac), since its enrichment at promoter regions follows the transcriptional activity (31, 32), and compared it with H3K27ac signal in whole muscle(33). In the non-sorted material 12550 and 12243 promoters with H3K27ac peaks were identified in soleus and EDL respectively, and 27% and 14% of these did not have enrichment after purification (Fig. 3A-F). For both muscles, gene ontology analysis of the promoters with signal only in nuclei from the whole tissue showed they were related mainly to the immune system and angiogenesis (Fig. 3 D -E and Table S1). Extending the analysis to the full epigenomes for whole muscle tissue and the purified myonuclei for the two muscle types, showed similar results with enrichment for angiogenesis and blood related functions (Fig. S3 A-D and Table S1).

**Fig. 3:**
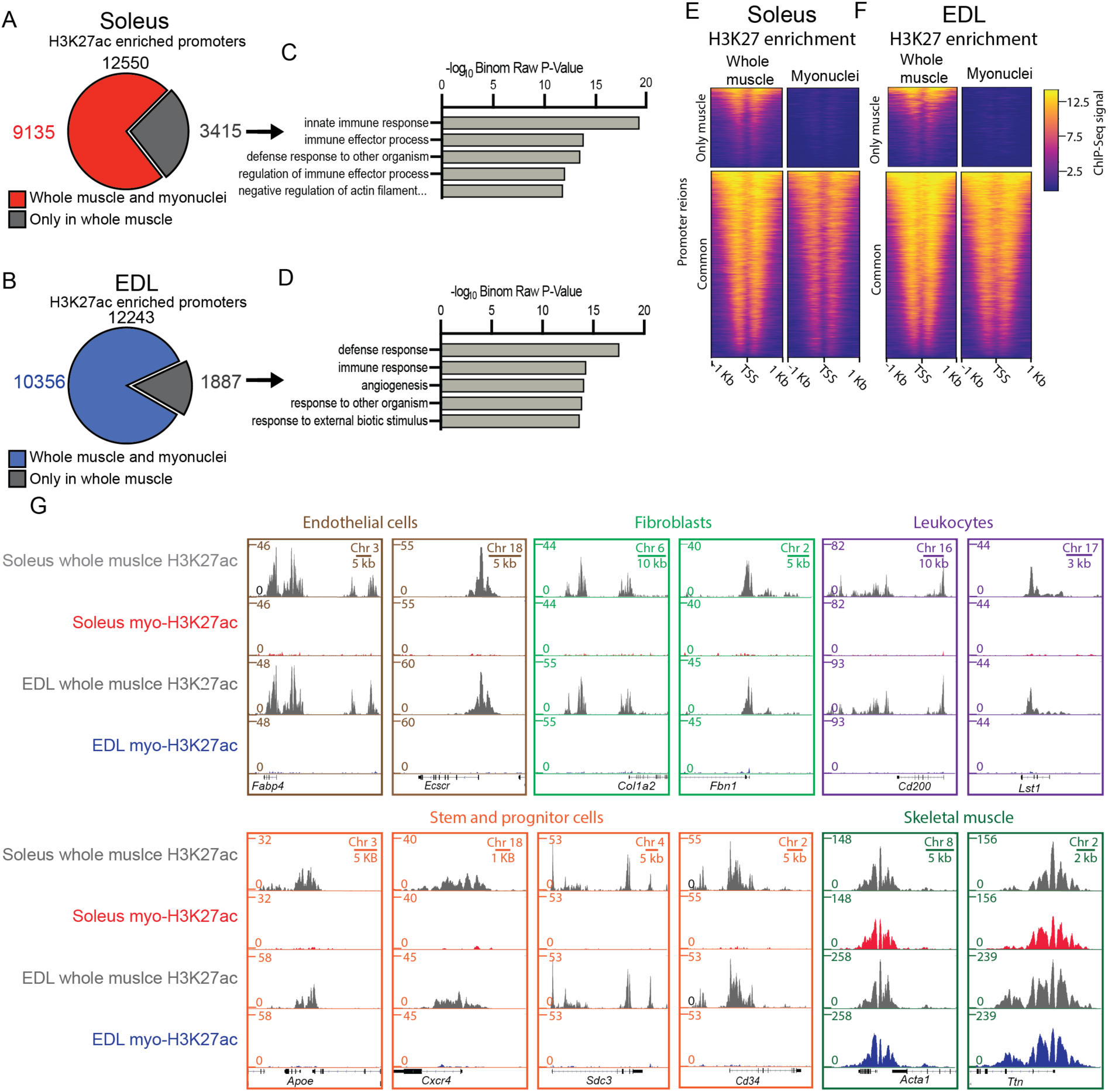
The epigenome of whole tissue is different from the myonuclei specific. A-B) Comparison of H3K27ac enrichment at gene promoter regions from whole muscle tissue and PCM1 isolated myonuclei for soleus and EDL respectively. C-D) Gene ontology enrichment analysis of promoter regions with H3K27ac enrichment specific for whole muscle tissue (grey). See Table S1 for full list. E-F) Heatmaps showing H3K27ac enrichment at the promoter regions in A-B -whole tissue and sorted myonuclei, in soleus and EDL respectively. G) H3K27ac enrichment in whole tissue and sorted myonuclei for loci used to define cellular populations.

The effects of purification was well illustrated by the observation that genes used to identify non-muscle cells were found to be enriched for H3K27ac in non-purified nuclei, but not after the purification, e.g. *Fatty Acid Binding Protein 4* (*FABP4*) and *Endothelial Cell-Specific Molecule 2* (*ECSCR*) which are both used as marks for endothelia cells (34-38). Concerning fibroblasts, the two connective tissue genes, *Collagen* (*COL1A2*) (34, 39-41) and *Fibrillin 1* (*FBN1*) (42-44) were found to be enriched before purification but not after. Similar observations were made for the two leucocyte markers *Leukocyte Specific Transcript 1* (*LST1*) and *CD200 Molecule* (*CD200*) which play a role in the immune defense (45, 46). The purified myonuclei did not appear to contain nuclei from satellite cells. Thus, markers common for satellite cells such as *Apolipoprotein E* (*APOE*) (34, 35) and *C-X-C Motif Chemokine Receptor 4* (*CXCR4*) (47, 48), *Syndecan-3* (*SDC3*) (49-52) and the general progenitor and stem cell marker *CD34* (53-56) was enriched in the total nuclei from the whole tissue, but not after purification of the myonuclei.

The loci for genes classically known to be a part of the skeletal muscle cell such as the skeletal specific version of actin *ACTA1* and *Titin* (*TTN*) displayed enrichment in both the full tissue and PCM1-sorted population (Fig. 3G).

### The epigenetic landscape in purified myonuclei from EDL and soleus reflects the differences in functional properties

To explore how the epigenome differed between the slow/oxidative soleus and fast/glycolytic EDL muscle cells, we focused on differentially enriched (DE) peaks between the two muscles using the histone modifications H3K4me3 and H3K27ac (Fold change (FC) >1.5 and a false discovery rate (FDR) < 1×10^−2^ and 1×10^−7^ for H3K4me3 and H3K27ac, respectively). We identified 719 H3K4me3 and 5309 H3K27ac DE peaks, corresponding to 7,6 % and 22,2 % of the sites, for each of these two histone modifications (Fig. 4 A-B). For H3K4me3 the majority of sites were localized around the promoter region, while for H3K27ac for most of the sites were found at distal regulatory enhancer regions (Fig. S4A-F). The higher number of DE peaks for H3K27ac suggesting that distal regulatory enhancers play important role in regulating the differences in function and phenotype, which is in line with previous observations of the epigenetic landscape between cell types from the same family (57, 58).

**Fig. 4:**
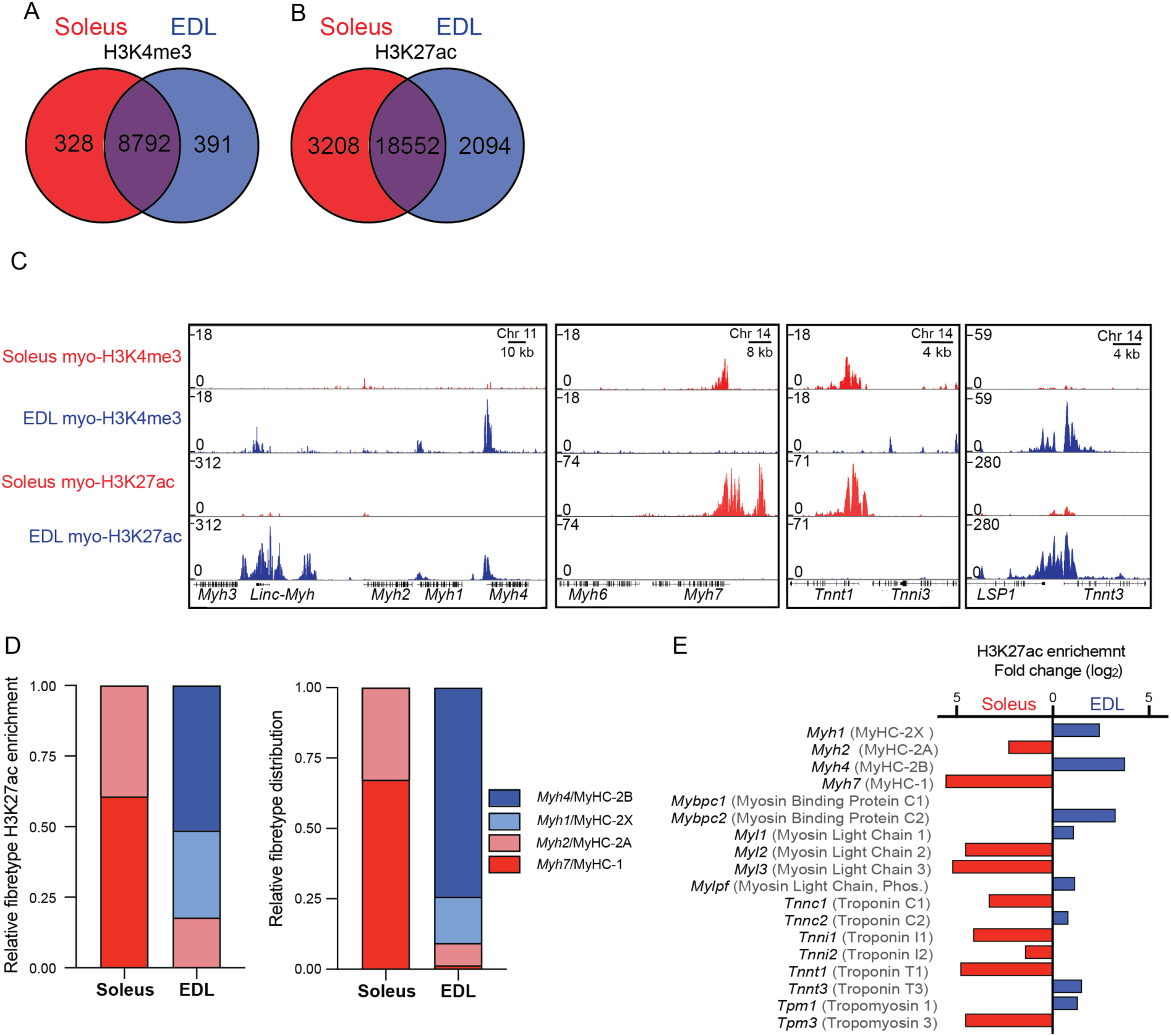
The differences in physiology are also present at the epigenetic leve1. A-B) Venn-diagrams showing number of peaks with similar or different enrichment for the histone marks H3K27ac and H3K4me3 in myonuclei. C) ChIP-Seq profiles of the myo-specific H3K4me3 and H3K27ac enrichment at the genomic loci containing the genes coding for the contractile proteins defining the muscle types: the main myosin types at the MyHC locus at chromosome 11 (*Myh1,-2,-4*) coding for the fast isoforms as well as the embryonically expressed *Myh3* and the non-coding RNA *Linc-Myh*, the slow *Myh7* gene at chromosome 14, and the Troponin genes the slow *Tnnt1* and the fast *Tnnt3* at chromosome 7. D) Relative H3K27ac enrichment for peaks overlapping the promoter regions for the four-principle myosin heavy chains (*Myh1,2,4 and -7*) and protein level for the corresponding proteins assessed by immunohistochemistry (MyHC1, MyHC-2A, -2B and -2X). E) Fold change differences in SOL/EDL in the H3K27ac enrichment in peaks overlapping the promoter region for the genes coding for phenotype defining contractile proteins. Gene names in black, protein names in grey. No significant difference in the enrichment for *Mybpc1*

#### The epigenome reflects fiber type specific differences in contractile properties

The most fundamental way of classifying muscle fiber types has been the particular MyHC gene expressed in the fiber, but the sarcomeres differ with respect to other contractile proteins as well (59, 60). These features were reflected in the epigenome (Fig. 4C). Thus, the slowest MyHC isoform *Myh7* (coding for MyHC-1) on the mouse chromosome 14 had a larger enrichment in soleus compared to EDL, while the opposite was true for the fastest form *Myh4* (MyHC-2B) that is clustered with the other type 2 heavy chains on the mouse chromosome 11 (Fig. 4C). When considering all the four limb muscle MyHCs, the H3K27ac enrichment roughly resembled the histological fiber type distribution of the two muscles that we have found previously (61) (Fig. 4E). Similar differences in the epigenetic enrichment were found at the loci coding for the other sarcomeric proteins, such as the myosin light chains (*Myl1-3*) (Fig. 4D). The fast isoform of the gene coding for the contraction regulator *Myosin Binding Protein C2 (Mybpc2)* was enriched in the EDL, while paradoxically gene coding for the slow isoform *Myosin Binding Protein C1* (*Mybpc1*), was not in the soleus, however this is in agreement with RNA and protein levels for the isoform (62-64)

As expected, the loci containing the genes encoding developmental isoforms of the sarcomeric proteins *Myh3* (MyHC-emb) and MYH8 (MyHC-neo) (Fig. 4C and Fig. S5) or isoforms specific for cardiac myocytes *Myh6* (MyHC-alpha) and *Troponin I3* (*Tnni3*/Troponin I), did not have any enrichment in these adult skeletal muscles (Fig. 4C).

#### The epigenome reflects fiber type specific differences the twitch speed

One of the major differences between different fiber types is the twitch duration, hence the EDL is a so called fast-twitch muscle and soleus a slow-twitch muscle. A major determinant of the twitch speed is the handling characteristics of Ca^2+^ (4, 60), and the expression level of relevant isoenzymes differs between different types of muscles (9, 41, 64, 65). Our data showed that this was also reflected at the epigenetic level such as for the, *ATPase Sarcoplasmic/Endoplasmic Reticulum Ca2+ Transporting* (*Atp2a*)*-1 and -2* (SERCA1 and -2), responsible for regulating the calcium levels in fast and slow muscle fibers respectively, *Casq1* and *-2* coding for the two calcium buffer proteins, *Calsequestrin 1* and *-2*, and *Pvalb* coding for the calcium binding protein Parvalbumin, highly specific for fast fibers (9) (Fig. 5).

**Fig. 5:**
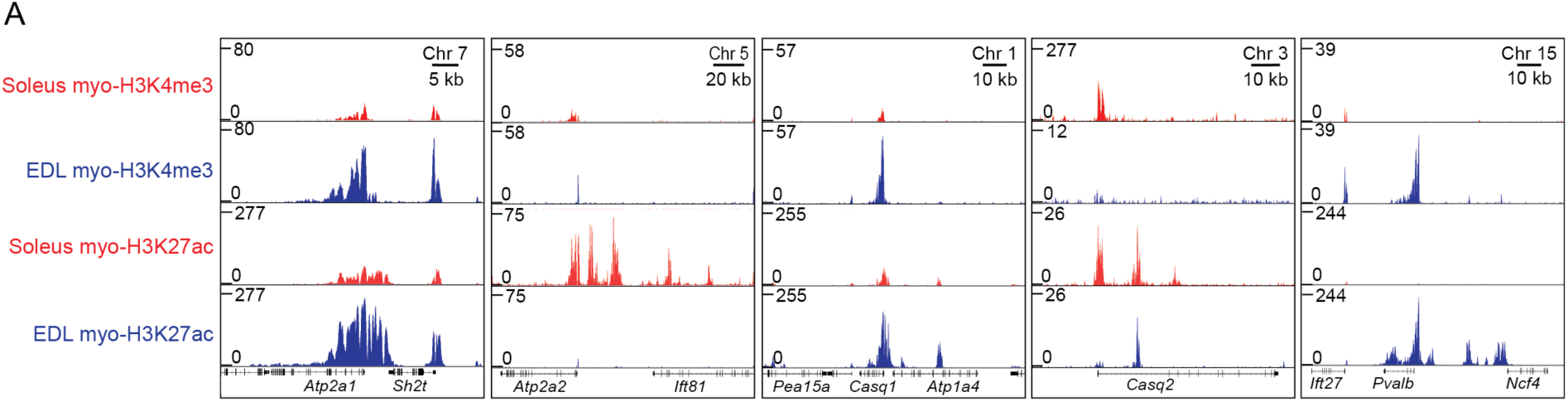
Genes involved in the calcium signaling have a diverse epigenome. A) ChIP-Seq profiles of the myo-specific H3K4me3 and H3K27ac enrichment at the genomic loci containing the genes coding for calcium signaling proteins: *ATPase Sarcoplasmic/Endoplasmic Reticulum Ca2+ Transporting (Atp2a1*,SERCA1) at chromosome 7, *Atp2a2* (SERCA2) at chromosome 5, *Calsequestrin 1* (*Casq1*) at chromosome 1, *Calsequestrin 2* (*Casq2*) at chromosome 3 and *Parvalbumin* (*Pvalb*) at chromosome 15, in soleus and EDL, respectively.

For the genes coding for contractile and calcium proteins, the differences in the epigenome were not only restricted to the promoter regions, but also the inter- and intragenic areas with H3K27ac enrichment, such as the enhancer located inside the MyHC cluster at chromosome 11 between the long non-coding RNA *Linc-Myh* and *Myh2* (MyHC-2A) promoting fast glycolytic phenotype (66), and upstream enhancer regions at the slow isoforms of *Myh7* (MyHC-1) and *Atp2a2* (SERCA2) genes (67) (Fig. 4B and Fig. 5). Furthermore, we also identified the recently discovered fast phenotype specific enhancer located in the locus coding for the *Pvalb* (Parvalbumin) gene (33). This indicates the complexity and detailed regulation of the genes involved in contraction and calcium handling in the different muscle types.

#### The epigenome reflects fiber type specific differences in metabolic properties

Ontology analysis of the H3K27ac DE peaks showed that the regions are associated with genes involved in defining the metabolic properties for the two muscle types, such as lipid metabolism and mitochondria for soleus and muscle contraction and metabolism of hexose (e.g. sugar) for EDL (Fig. 6A-B and Table S2). Interestingly, a similar analysis carried out on whole tissue, identified fewer DE regions and only non-specific general functions (Fig. S4 G-H and Table S2). Again, underscoring the importance of sorting out the relevant nuclei from heterogenous tissues.

**Fig. 6.**
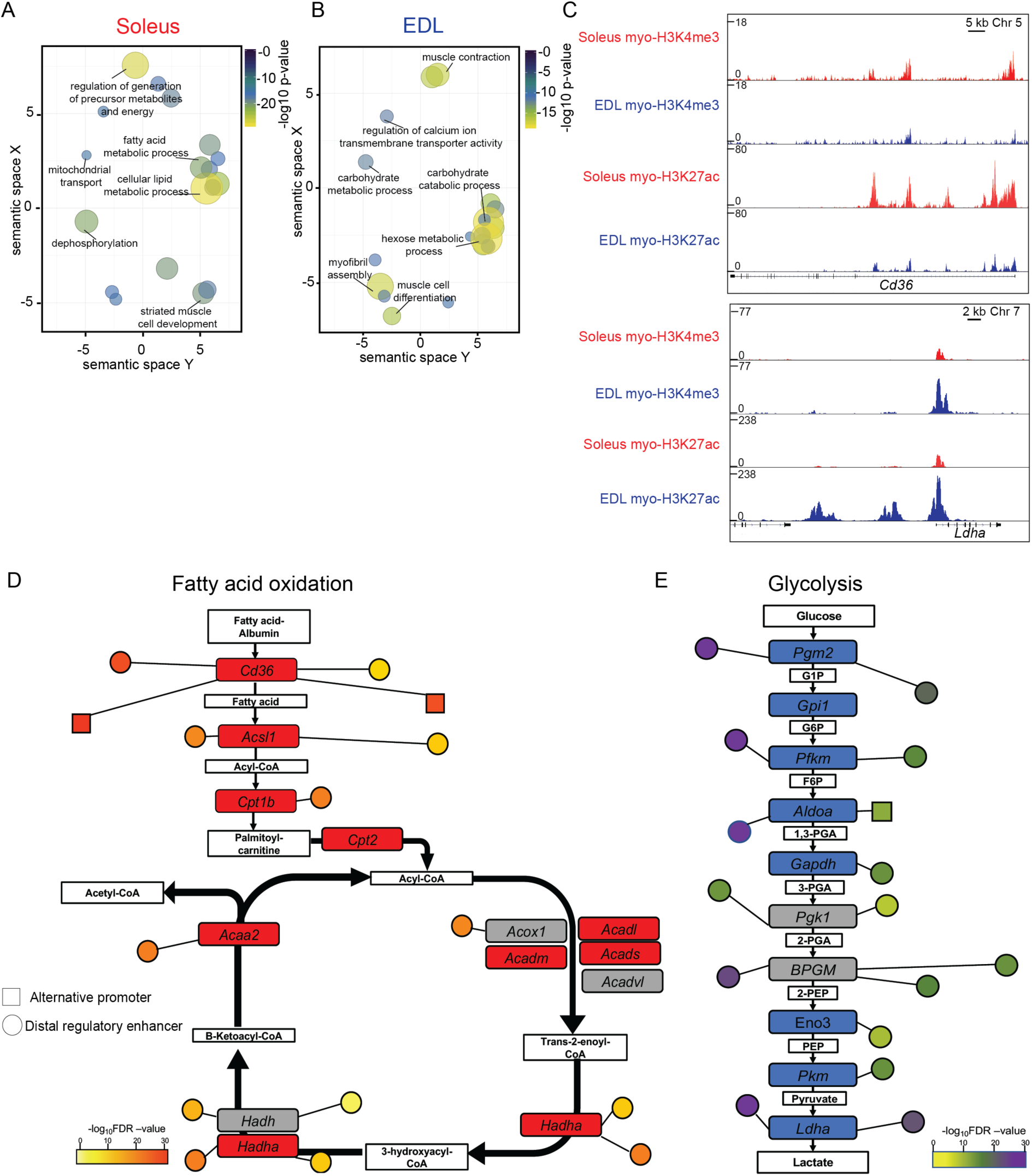
The epigenomes of the muscle extremes are enriched for different metabolic profiles. A-B) Plot of enriched gene ontologies for myo-specific differently H3K27ac peaks enrichment in soleus and EDL, respectively. The ontologies are shown after redundancy reduction with ReViGO by functional similarity. Terms are clustered in the semantic space by their similarity, without intrinsic meaning to the semantic space units. Selected terms for each cluster are shown, see Table S2 for full list. Ontologies identified with GREAT using closest promoter. C) ChIP-Seq profiles of the myo-specific H3K4me3 and H3K27ac enrichment in soleus and EDL, for two loci with differently enrichment, encoding *Fatty acid translocase* (*Cd36/Fat*) at chromosome 5 and *Lactate dehydrogenase A* (*Ldha*) at chromosome 7, respectively. D-E) Schematic representation showing the genes encoding the enzymes involved in the fatty-acid oxidation and glycolysis, respectively. Genes with a differently enriched H3K27ac promoter colored in red and blue for soleus and EDL, respectively. Grey denoted no difference in enrichment. Differently enriched alternative promoters and enhancer regions are shown as nodes with relative distance to the primary promoter region. Color code denotes the FDR significance level of the differential enrichment between the two muscle types.

To further gain insight into how the epigenome adds to differences between the two types of muscles, we examined the differences in biological and signaling pathways using the dynamic H3K27ac mark. The analysis identified that key factors and enzymes responsible for the metabolism and muscle size and -function have a different H3K27ac enrichment, both at promoters and distal regulatory enhances regions; such as Peroxisome proliferator-activated receptors (PPAR)- and the Fatty acid degradation pathways in soleus and the glycolysis and mitogen-activated protein kinase (MAPK) pathways in EDL (Table S3). Detailed analysis of the epigenetic landscape for two of the primary energy generating pathways, the fatty acid oxidation in soleus and glycolysis in EDL identified a complex epigenetic environment with several promoters and enhancers with a differently enrichment for the genes involved in the pathways (Fig. 6 C-E). For both the fatty acid oxidation pathway and glycolysis none of the involved genes were associated with DE regions for the opposite muscle type e.g. glycolysis enriched in soleus and fatty acid oxidation in EDL.

### The epigenome indicates that the phenotype of fast/glycolytic and slow/oxidative muscles are determined by different regulatory networks

We sought to unravel the gene regulatory programs leading to the fast/glycolytic and slow/ oxidative phenotypes respectively, by using the myonuclei-specific DE H3K27ac regions in combination with DNase I hypersensitivity data for whole skeletal muscle (68, 69), which with a high resolution identifies where transcription factors and other proteins are bound to the chromatin (70). Motif enrichment analysis of the DE regions identified two distinct groups of transcriptional regulators for the two muscles. In soleus the SRY-Box Transcription Factor (SOX) and Nuclear Factor Of Activated T Cells (NFAT) motifs were the most overrepresented, while in EDL it was the E-boxes and Sine Oculis Homeobox Homolog (SIX) motifs (Table S4).

To consolidate and specify the analysis, we used the transcription factors from the motif enrichment analysis with the most significantly enriched promoter region and high quality motifs for each muscle. This revealed two distinct groups of transcription factors one for each muscle type; for soleus the factors such as SOX6, NFATC2, Myocyte Enhancer Factor 2C (MEF2C) and Peroxisome Proliferator-Activated Receptor Alpha (PPARA) were identified while for EDL factors such as Myogenic Differentiation1 (MYOD1), SIX1 and Zinc Finger and BTB Domain Containing 14 (ZBTB14) (Table S4). The transcription factors were identified to have 4530 and 4371 predicted binding sites connected to 464 and 392 DE promoter regions for soleus and EDL, respectively. Functional analysis of the genes connected to the transcription factors revealed that they are associated with central differences in the functional phenotype between the two muscles (Table S5). Combining the information about factors, genes and functions revealed distinct regulatory networks for the two muscles (Fig. 7). Though different, they also share similarities, such as regulation of muscle contraction, but with a different set of factors and genes involved (Fig. S6).

**Fig. 7:**
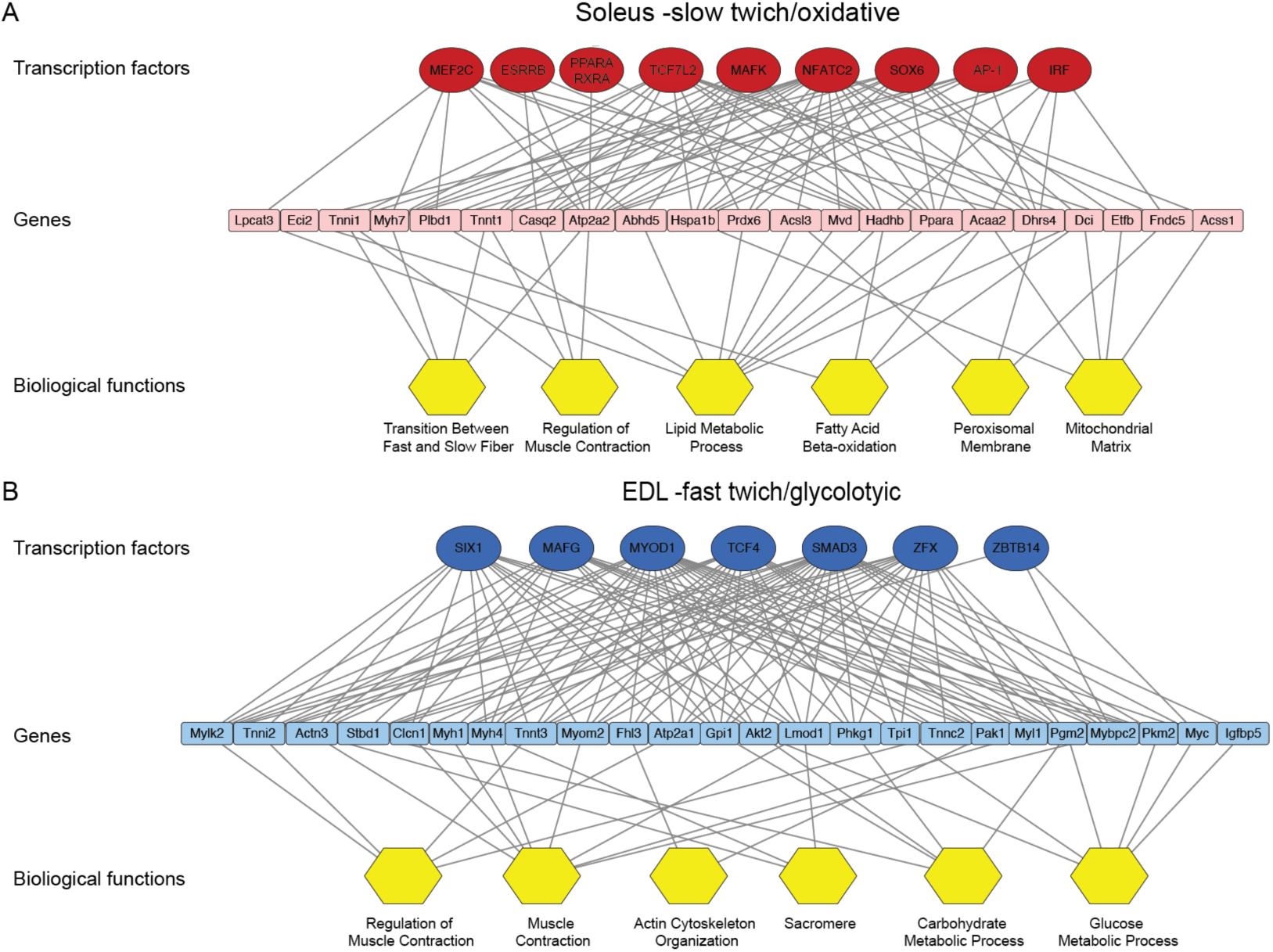
The muscle extremes are enriched for different regulatory networks. Transcriptional regulatory network for soleus (A) and EDL (B). Top row constitutes of transcription factors with motif enriched in DE H3K27ac peaks, the middle of associated genes with a DE promoter (< 100 kb). The last row shows biological functions assigned to the genes. Edges between transcription factor and genes represent identification of predicted binding site for the respective transcription factor site and the gene it is connected to. Edges between genes and biological function denotes the functional properties assigned to the gene. Transcription factor motifs were identified with AME using high-quality motifs from human and mouse FDR < 1×10^−5.^ For factors with similar motifs, the factor with the most significantly enriched promoter in the respective muscle type was used. Biological functions represent functional annotation clusters associated with the differential enriched genes (Clusters identified with David using an enrichment score >2.2 for the clusters and a p-value and fold change for the ontologies at < 5×10^−2^ and >3, respectively). Network visualized with Cytoscape. For the sake of clarity, only selected genes, with their most significant edge to each transcription factor, and terms are shown. See Table S4 for the full names of transcription factor of factors and motifs and Table S5 for functional annotation clusters and genes.

## Discussion

We show that PCM1 selectively marks mature myonuclei in skeletal muscle, and that this marker can be used to isolate the myonuclei from wild type fresh and frozen muscle samples. This isolation method does not involve any enzymatic digestions or other treatments at elevated temperatures, thereby preserving the native state of the epigenome. Furthermore, the method relies on magnetic sorting, which simplifies the procedure. The sorting is important since only less than 60% of the cell nuclei in muscle tissue are myonuclei (22, 24), and the sorting excluded the large fraction of nuclei from satellite cells and stroma cells such as connective tissue and vascular cells. The epigenetic signature of individual genes is highly cell-specific Studies of other heterogenous tissues have also concluded that accounting for cellular heterogeneity is critical in epigenome-wide association studies (25-27). Mixed cell populations are prone to increase the number of false negatives and positives (32, 71). This point has also been addressed in skeletal muscle by mechanically isolating single fibers preceding transcriptome (41) and methylome (72); reducing the signal arising from non-myogenic cells residing in the tissue.

When we compared myonuclei from the two extreme muscles soleus and EDL we found that they had a large part of the epigenome in common. Important differences were found in connection to genes coding for proteins involved in contraction, calcium handling and metabolic pathways; reflecting differences at the transcriptional and protein level between different fiber types (41,64,73-75).

In addition, we found differences in distal enhancers at the loci reflecting different regulatory network in the fast and the slow muscle. The two MyHC loci provide interesting examples of differences in such regiona lregulation. In both mice and man, the slow type 1 *Myh7* (MyHC-1) gene is located on chromosome 14 head to tail with the cardiac isoform *Myh6* (MyHC-1alpha), while the fast type 2 isoforms are clustered on a 300-600 kb segment in the order *Myh4, Myh2* and *Myh4*, reflecting the order of increasing of shortening velocity, MyHC-2A-2X-2B) on the mouse chromosome 11 (in humans 17) (76). Interestingly, during exercise, activity-induced changes are relatively easily obtained within the different type 2 MyHCs of the gene cluster, while 1⇔2 transformation is harder, and cannot be obtained, except under extreme conditions (7).

In the fast locus an intergenic SIX1 responsive enhancer between *Myh3* (MyHC-emb) and non-coding RNA *Linc-Myh* are located. It drives the expression of the fast myosin through higher order chromatin interactions and locks the muscle fiber in fast-glycolytic phenotype (66). Furthermore, the enhancer regulates the *Linc-Myh*, that additional play a role in maintaining the fast phenotype, presumable through trans-induced mechanisms (66). This area which is one of the most differently enriched in our data for the glycolytic EDL, revealed several additional enhancer regions upstream and downstream of the *Linc-Myh*, further adding to the complexity of the locus. In muscle cell culture this area has been identified to be a part of a special group of enhancers known as super-enhancers (77). This group of enhancers are characterized by spanning several kb and being important for defining the identity of the cell (78-80). As one of the most enriched area in EDL, it suggests that the super-enhancer also plays a role in the adult glycolytic fiber *in vivo* and might stabilize fiber types and be an obstacle to transformations.

For the regulatory network several of the transcription factors are well known regulators of skeletal muscle phenotype and function, such as NFAT, SOX, SIX and MYOD1 (4, 60, 81). Interestingly our enriched factors are largely in agreement with a recent study, using an alternative approach to investigate the epigenetic regulatory environment in slow and fast muscles at the single nuclei level (82).

Other factors we identified, such as MAF BZIP Transcription Factors (MAF) and ZBTB14, is just recently described to have a role in muscle regulation at the gene level (82, 83). The latter is identified to be involved in regulation of metabolism, a function also assigned to the factor in our regulatory network. More specifically, it is found to be a negative regulator of the myokines *Interleukin 6* (*IL6*) and *Leukemia inhibitory factor* (*LIF*) by binding to the promoter regions of the two genes (83). We identified two additional factors, largely unexplored in skeletal muscle namely the Zinc Finger Protein X-Linked (ZFX) in EDL and Transcription Factor 7 Like (TCF7L2) in soleus, but both has previously been suggested to be a part of skeletal muscle specific regulatory networks (64, 84). On the molecular level the ZFX is identified to facilitate transcription through binding in the promoter regions just downstream of the transcription start sites of target genes (85). In the case of TCF7L2, it is found to regulate the expression of the slow *Myh7* (MyHC-1) (86) and modulate the chromatin environment in oxidative muscle tissue (87).

For the soleus it is notable that the transcription factors MEF2C, PPARA, SOX6 and NFATC2 are identified as they are known to influence muscle in the slow/oxidative direction (60, 88-91). MEF2C is previously identified to be important for slow fiber development and for the energy homeostasis in these (92, 93).

Interestingly, MEF2C and its family members are known to interact with co-regulators that are capable in altering the epigenetic environment such as histone acetyl transferase P300 (94, 95) and lysine methyltransferase Myeloid/lymphoid or Mixed-Lineage Leukemia 4 (MLL4). In the latter case, the transcription factor together with its co-regulator establish an open chromatin environment at active enhancers that enforces an oxidative/slow muscle phenotype e. g. by regulating sarcomeric slow genes such as *Myh7* (MyHC-1) and *Tnnc1* (Troponin C1) through enhancers located upstream of their respective promoters (67). MEF2C is also participating in muscle activity-related calcium regulation, both by being activated by downstream signaling pathways such as MAPK and calmodulin-dependent protein kinases (CaMK) (96-98), but also through regulation of the genes participating in the signaling cascades (99-101). From the cardiac tissue the transcription factor is found to regulate the expression of the slow sarcoplasmic calcium pump, *Atp2a2* (SERCA2), through an enhancer region (102), a connection also identified in our soleus network. This points towards an interesting conserved regulatory mechanism between heart and skeletal muscle.

The transcription factors PPARA and Estrogen Related Receptor Beta (ESRRB) are mediating fatty acid degradation and oxidative metabolism (73, 103, 104). Similar roles are known for additional members of their respective families (105, 106). In the case of PPARA, it is furthermore reported to control the expression of metabolic genes in response to changes in the environment through distal enhancers (107) and to be involved in regulation of contraction (108, 109).

For EDL it was notable that SIX1 and MYOD1 appeared since there is evidence that they drive the phenotype in the fast/glycolytic direction (4, 110, 111). The SIX1 transcription factor is reported to be an important regulator of key genes defining the fast-glycolytic phenotype where it regulates genes mediating glycolysis (e.g. *Aldoa* (Aldolase A), *Pfkm* (Phosphofructokinase) and *Eno3* (Enolase 3) and calcium homeostasis (*Pvalb* (Parvalbumin) (112, 113) in addition to regulating the non-coding RNA *Linc-Myh* and its enhancer region (113). MYOD1 is found to be important for correct glycolytic fiber type specification and composition (111, 114). On the molecular level MYOD1 is known as a master regulator and is proposed to work as an organizer of enhancers (115, 116) In line with this it is known to recruit histone modifiers and co-regulators p300 and its homologue CBP to target sites (94, 117, 118).

A recent muscle cell differentiation study in cell culture, MYOD1 were identified to take part in the regulation of the super-enhancer inside the fast MyHC locus covering the *Linc-Myh* and its enhancer (119). Pointing towards an interesting collaboration at the locus between MYOD1 as a master regulator and the more specific role of SIX1 in locking the expression of the fast myosin genes (66) thereby promoting the glycolytic phenotype.

In summary we have shown that PCM1 can be used to isolate the myonuclei from the complex whole muscle tissue and that in order to get a clear and faithful representation of the whole epigenome, such sorting should be performed prior to analysis of nuclei from the complex muscle tissue. Comparison of the myo-specific epigenomes of the two muscle extremes soleus and EDL show as expected that the majority of the epigenome in common. The parts that differed were regulatory elements and genes related to contraction speed, twitch duration, and metabolism. In addition, our epigenetic analysis revealed that the two muscles have distinct regulatory networks associated with genes defining the disparate phenotype.

## Materials and Methods

### Materials and Methods

#### Animal procedures

Female NMRI and (28-31 gram) and Sprague Dawley rats weighing 210-260 g were used. Animals were kept at the animal facility at the Department of Biosciences or Department of Medicine, University of Oslo. Animals were housed with a 12 h light/dark cycle with ad libitum access to food and water. Before surgery, animals were sedated with 2 % isoflurane (506949, Forene, Abbot) in the air. Following deep anesthesia, the hind limb muscles extensor digitorum longus (EDL), soleus and tibialis anterior (TA) were surgically removed and directly frozen in liquid nitrogen and transferred to cryotubes before being stored at -80 °C. Animals were sacrificed by cervical dislocation while under deep anesthesia. The research was conducted in accordance with the Norwegian Animal Welfare Act of 20th December 1974. The Norwegian Animal Research Committee approved all experiments before initiation. The Norwegian Animal Research Authority provided governance to ensure that facilities and experiments were in accordance with the Act, National Regulations of January 15th, 1996, and the European Convention for the Protection of Vertebrate Animals Used for Experimental and Other Scientific Purposes of March 18th, 1986.

The ACTA1^rtTA^; TRE^H2B-GFP^ mice were kindly provided by Dr. John McCarthy (University of Kentucky)(30). To induce H2B-GFP expression, 2-month old mice were fed chow supplemented with Dox (625ppm) for 1 week. The Dox chow was purchased from TestDiet (54057). All procedures with the ACTA1^rtTA^; TRE^H2B-GFP^ mice were approved by the Cincinnati Children’s Hospital Medical Center’s Institutional Animal Care and Use Committee.

#### Immunohistochemistry and imaging

For cross section tissue sectioning, immunohistochemical staining and subsequent image processing were performed as previously described (24).

For single fiber imaging; EDL muscles from the ACTA1^rtTA^; TRE^H2B-GFP^ mice were collected and incubated in high-glucose DMEM (HyClone Laboratories) with 0.3% collagenase type I (Sigma-Aldrich) at 37°C for 40 minutes, then washed with PBS. Muscles were gently triturated using a glass pipette to loosen digested myofibers until they shed from muscle and fixed overnight in 1% paraformaldehyde. Fibers were then put in staining buffer (0.6% Igepal-CA-630, 5% BSA and 1% goat serum in PBS) for 1h at room temperature, followed by staining with an antibody against PCM1 (1:1000, HPA023370, Sigma Aldrich) in staining buffer over night at 4 °C with gentle agitation. Next day fibers were washed (3x 30 min in staining buffer), incubated for 3 hours with secondary antibody (Abcam #150083) at 4 °C with gentle agitation, washed 3x 30 min in staining buffer and mounted with Fluoromount-G with DAPI (SouthernBiotech #0100-20). For Figure 1D-E, fibers were imaged on a Nikon A1 R confocal system through NIS-Elements AR software (ver.5.10.01) with a 60x oil immersion objective with NA of 1.4, using a Galvano scanner. For Figure S1, fibers were imaged on a Nikon Eclipse Ti inverted microscope with a 20x air immersion objective.

#### Myonuclear Isolation

All samples and buffers were kept on ice during the procedure. The muscle was removed from the -80 °C freezer and transferred to -20 °C where it was minced into pieces of 1-2 mm before being transferred to gentleMACS M-tubes (Miltenyi Biotec) tubes kept on ice. The samples were homogenized in 5 ml of lysis buffer (10 mM Tric-HCl pH 8.0 (cold adjusted), 5 mM CaCl_2_, 3 mM MgAc, 2 mM EDTA, 0,5 mM EGTA) on a gentleMACS Dissociator (Miltenyi Biotec) using the default homogenization program *protein_01*. All buffers were supplemented with 5 mM Na-butyrate (Sigma Aldrich), 5 mM PMSF (Sigma Aldrich) and 1x protease inhibitor (Complete Protease Inhibitor Cocktail, Roche) immediately before use. Following homogenization, the lysate was diluted in 5 ml lysis buffer containing 0.4 % Triton-X100. Samples were mixed 10 times with a 1 ml pipette before being filtered through a 100 *μ*m and 30 *μ*m strainers (Falcon, Sigma Aldrich) to remove large aggregates. The homogenates were transferred to a 15 ml tube and centrifuged at 1000 g for 5 minutes in a centrifuge with a swing-out rotor at 5 °C. The pellet was resuspended in 1 ml nuclei staining buffer (5 % BSA wt/vol, 0.2 % IGEPAL–CA630, 1 mM EDTA in Dulbecco’s Phosphate Buffered Saline (DPBS) pH 7.4) (120) containing an antibody against PCM1 (1:1000, HPA023370, Sigma Aldrich). Samples were incubated using a tumbler (40 rpm) for 1 hour at 6 °C. Then centrifuged for 5 minutes at 800 g at 5°C and the pellet was resuspended in 500 *μ*l sorting buffer (1 % BSA wt/vol, 2 % skimmed milk powder (Sigma Aldrich), 1 mM EDTA in PBS pH 7.4) and spun down at 600 g at 5°C for 5 minutes at slow acceleration settings. The pellets were resuspended in 100 *μ*l beads buffer consisting of 80 *μ*l sorting buffer with 20 *μ*l secondary Anti-rabbit IgG MicroBeads (Miltenyi Biotec) and incubated for 15 min in at 4 °C sorting buffer (900 *μ*l). After incubation, the samples were centrifuged as above and wash once with Sorting buffer before resuspended in 1 ml of the same buffer and incubated for 5 minutes at a tumbler at 40 rpm at 6 °C. After incubation, the sample was carefully applied to a M column (Miltenyi Biotec), washed 3 times with 1 ml Sorting buffer and eluted with 1 ml Sorting buffer after removal from the magnet. The eluate was applied to another M column, washed again and eluted in 1 ml elution buffer (DPBS with 1 mM EDTA). The efficiency of sorting was determined by staining aliquots before and after myonuclear enrichment.

#### Chromatin immunoprecipitation

All buffers were supplemented with 5 mM Na-butyrate (Sigma Aldrich), 5 mM PMSF (Sigma Aldrich) and 1x protease inhibitor (Complete Protease Inhibitor Cocktail, Roche) immediately before use. Nuclei, approximately 300 000, were fixated with 1 % freshly prepared formaldehyde (Fisher Scientific) in DPBS for 2 min at room temperature. Reaction was stopped by adding glycine to a final concentration of 125 mM. After 5 minutes of incubation at RT, nuclei were transferred to ice and washed two times with ice cold DPBS and lysed in 130 *μ*l sonication buffer (50 mM Tris-HCl pH 8.0 (cold adjusted), 1 mM EDTA, 0.1 % wt/vol SDS). Samples were sonicated for 8 minutes (30 sec on/off cycles) using a Bioruptor Pico (Diagnode) yielding on average DNA fragments of 200 -300 bp. Lysate was diluted in an equal volume of 2x RIPA buffer (20 mM Tris-HCl pH 8.0 (cold adjusted), 280 mM NaCl, 1 mM EDTA, 1 mM EGTA, 2 % Triton X-100, 0.2 % Na-deoxycholate, 0.1 % SDS), and adjusted to 800 *μ*l with RIPA buffer (10 mM Tris-HCl pH 7.5 (cold adjusted), 140 mM NaCl, 1 mM EDTA, 0.5 mM EGTA, 1 % Triton X-100, 0.1 % Na-deoxycholate, 0.1 % SDS). Samples were centrifuged at 12 000 g for 10 minutes and the supernatant was transferred to new tubes and precleared with 10 *μ*l Protein A Dynabeads (Invitrogen) for 1 hour. For each sample, ChIP assays were performed in parallel with antibodies directed against H3K27ac (C15410196, Diagnode) and H3K4me3 (C15410003, Diagnode) or IgG (Kch-504-250, Diagnode). Antibodies (2 *μ*g per sample) were incubated 10 *μ*l to Protein A Dynabeads for two hours on a rotator at 40 rpm at 6 °C. Beads were transferred to PCR-tubes and captured with a magnetic rack followed by addition of 150 *μ*l ChIP ready chromatin. Samples were incubated at 40 rpm on a tube rotator overnight at 40 rpm at 6 °C. Next morning, the supernatant was removed and the beads was washed with 100 *μ*l RIPA-buffer, followed by a washing step with high salt RIPA (20 mM Tris-HCl (cold adjusted) pH 8.0, 500 mM NaCl, 1 mM EDTA, 1 mM EGTA, 2% Triton X-100, 0.2 % Na-deoxycholate, 0.1 % SDS) and a washing step with LiCl buffer (20 mM Tris-HCl pH 8.0 (cold adjusted), 250 mM LiCl, 1 mM EDTA, 1 mM EGTA, 2 % Triton X-100, 0.2 % Na-deoxycholate, 0.1% SDS) and once with TE-buffer (20 mM Tris-HCl pH 8.0). All washing steps were being performed for 5 minutes using a rotor at 40 rpm and at 6 °C, with the exception of the TE-buffer washing steps which was carried out at room temperature. After the final washing step, the beads were resuspended in 96 *μ*l elution buffer (20 mM Tris-HCl pH 7.5, 5 mM EDTA, 50 mM NaCl, 1 % SDS) and incubated on heating block at 37°C. After one hour, 1 *μ*l Proteinase K (20 µg/ul) was added to each sample and incubated for one hour at 50°C followed by 4 hours at 68°C on thermoshaker at 1200 rpm. ChIP DNA was purified with Zymo ChIP DNA Clean & Concentrator kit in 10 *μ*l (Zymo Research).

#### Flow cytometry

For flow cytometry analysis nuclei were isolated as described above. For the ACTA1^rtTA^; TRE^H2B-GFP^ TA muscles were excised, minced with a razor, and homogenized in PBS containing 0.25M sucrose and 1% BSA using an Ultra-Turrax T25. The homogenate was then incubated for 5 minutes with addition of Triton-X100 to a final concentration of 0.36 % at 4°C. Samples were filtered through a 100 µm strainer, then filtered again through a 40µm strainer. The nuclei pellet was collected after centrifugation (3000?×g for 5?minutes at 4°C), and resuspended in sorting buffer (2 % BSA/ PBS). PCM1 (HPA023370, Sigma Aldrich) staining was performed on ice for 1 hour (1:1000). After washing, samples were stained with the secondary antibody Alexa Fluro 647 anti-rabbit IgG (A32795, Invitrogen)(1:1000) for 30 minutes in combination with Hoechst 33342 (Life Technologies) added 1:5000. Stained samples were analysed on a BD FACSCanto II or a BD LSRII. Data analysis was performed utilizing BD FACS Diva, FlowJo and Kaluza.

#### Next-generation sequencing

Libraries for next-generation sequencing (NGS) were created with the Swift Accel-NGS 2S Plus DNA Library Kit (Swift Biosciences) following the recommendations from the manufacturer with the exception that the ratio of beads and PEG was 1.5 and 1.3 in step Repair step I and II, respectively and amplified with 12 PCR cycles. The libraries were sequenced on a HiSeq 2500 40 bp paired-end, Illumina at the Norwegian Sequencing Centre.

#### Bioinformatics analysis

The ChIP-Seq reads were trimmed with TrimGalore (121) and mapped to the mouse genome (mm9) using BWA (122). Duplicated and low-quality reads were removed (q >10) using Samtools (123). Enriched regions were identified with PePr using the three biological replicates for each histone modification over the respective input samples using the following parameters --sharp for peak calling and a p-value threshold on 0.05 (124). Differently enriched regions were identified using PePr --diff with the biological replicates over their respective input for each sample and using threshold on foldchange 1.5 or above and a FDR-value on 1×10^−5^ for H3K4me3 and 1×10^−7^ for H3K27ac or below in addition to intersect with peaks identified in their respective samples.

For gene annotation UCSC Known Genes (mm9) with an ensemble id were used. Promoters were defined as upstream of 750 bp and downstream of 250 bp of transcription start site (TSS). For visualization samples were normalized to input and BigWig files were created as described in (125). Heatmap of ChIP-Seq signals were created with deepTools version 3.3.0 (126) Gene ontology analyzes of DE peaks were performed with GREAT v. 4.04 (127) using single closets gene while pathway analysis was conducted with David v. 6.8 (128, 129). Redundancy reduction was performed with ReViGO (130).

Regulatory networks were visualized with Cytoscape (131). DNase I hypersensitive enrichment data for mouse skeletal muscle were obtained from (68, 69). For the analysis, only middle point of DNase I enriched areas -/+ 100 bp inside the myo-specific H3K27ac enriched areas for Soleus and EDL were used. Motif enrichment analysis was conducted with AME Version 5.0.4 (132) using high quality motifs from human and mouse and threshold on FDR < 1×10 ^−5^. For factors with similar motif, the factor with the most significant H3K27ac enrichment at the promoter region (FDR < 1×10 ^-15^) was included. Functional annotation clustering of enriched promoters was conducted with David v. 6.8 (128, 129).

Data have been made publicly available under GEO (accession number GSExxxxxx)

## ACKNOWLEDGEMENTS

We would like to thank Inga Juvkam for technical assistance. Bioinformatics analyses were performed at the Abel Cluster (project nn9540k), owned by the University of Oslo and the Norwegian metacenter for High-Performance Computing (NOTUR).

