## Supplementary figures S1-S5 for "The epigenetic landscape in purified myonuclei from fast and slow muscles"

### Supplementary figures Bengtsen et al.

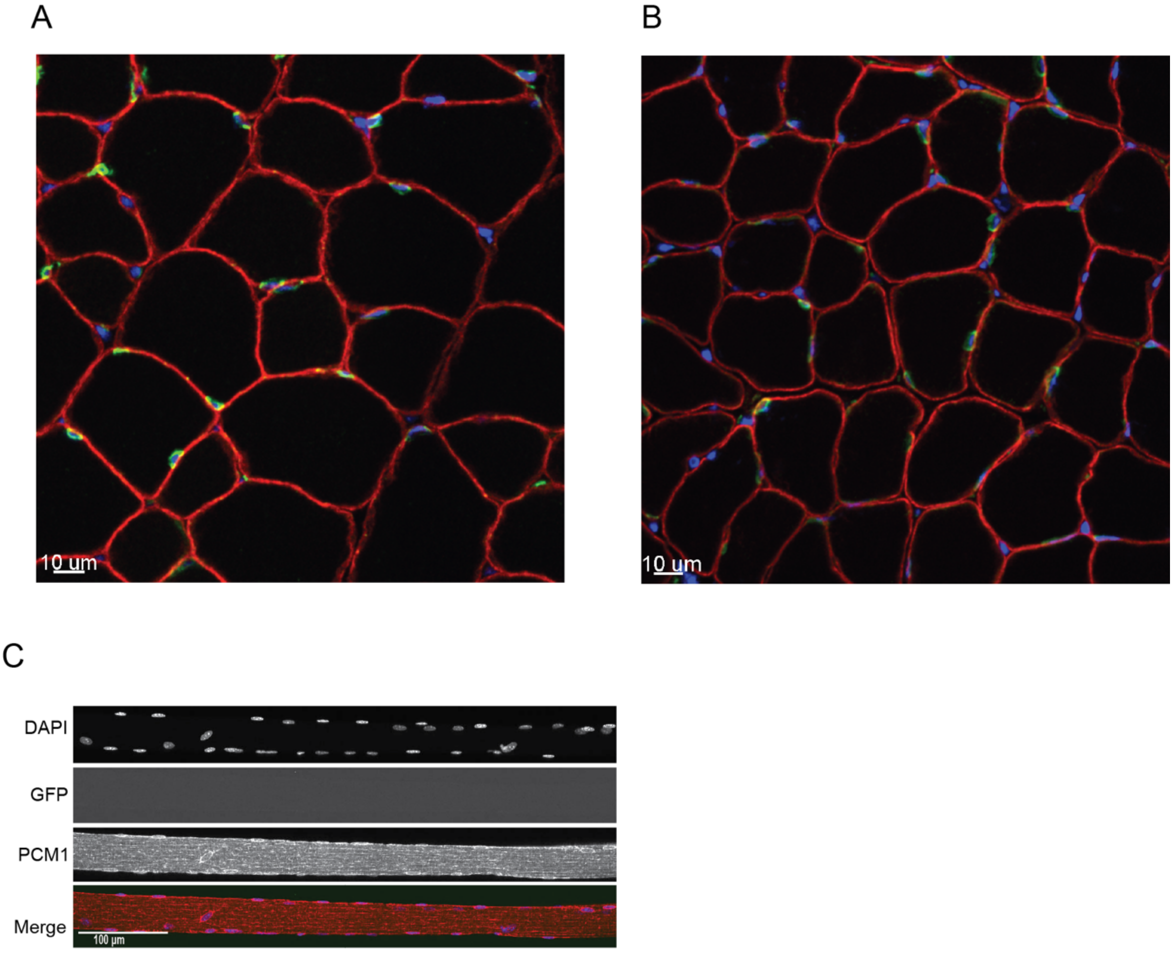


**Fig. S1: PCM1 selectively labels the myonuclei on muscle cross-sections and isolated nuclei.** A-B) Cross-section of EDL and soleus muscles stained with antibody against PCM1 (green), dystrophin (red) marking the boundary of the myofibers. Counterstained with DAPI visualize DNA (Blue). Scale bar 10 $\mu$m. C) Max intensity projection of a single fiber from a wild-type control mouse corresponding to the GFP and PCM1 co-localization in Figure 1. Scale bar 100 $\mu$m.

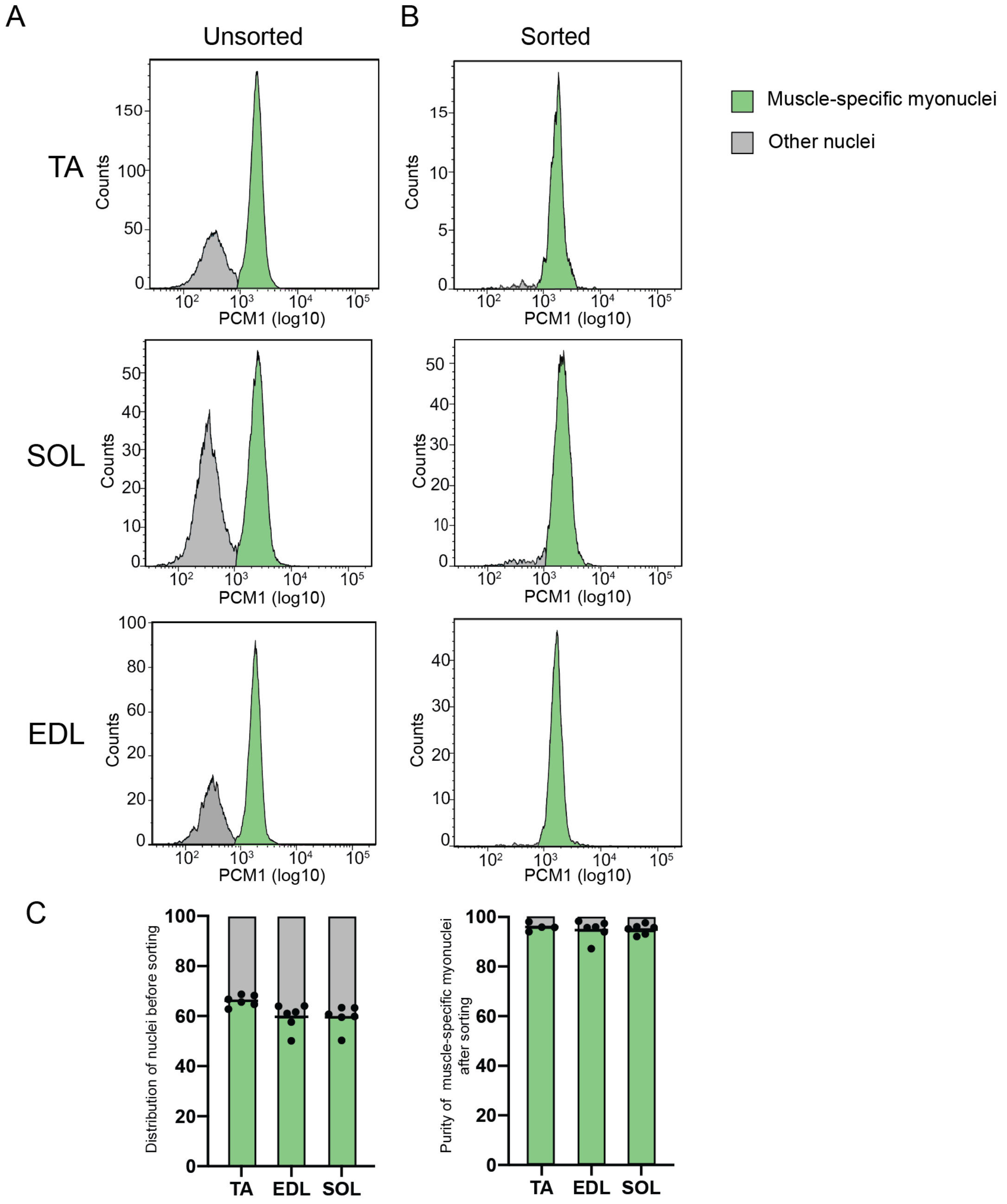


**Fig. S2 Analysis of myonuclei distribution and isolation in skeletal muscles from rat.** A) representative histograms of rat nuclei distribution for the three muscles TA, EDL and Soleus by flow cytometry. B) Magnetic sorting efficiency of myonuclei for the three muscles. C) quantification of nuclei distribution and sorting efficiency after sorting (n=4-6).

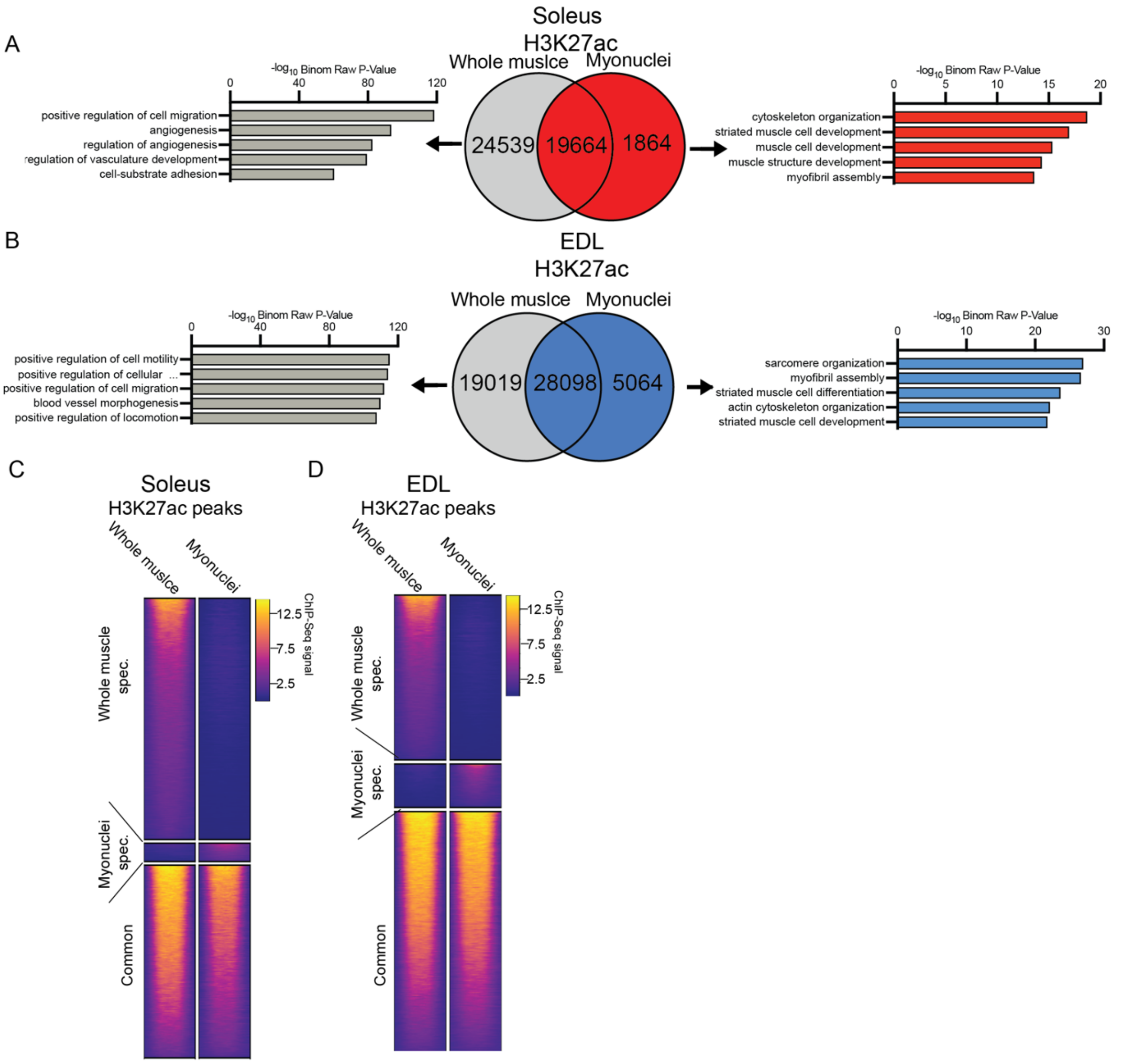


**Fig. S3: Differences in the epigenome between whole muscle and sorted myonuclei.** A-B) Venn diagram unique and common H3K27ac peaks between soleus and EDL whole muscle and myonuclei, respectively. The five most enriched gene ontologies for the unique peaks are shown to the left (whole tissue) and right (myonuclei) For full list of ontologies see Table S2 Gene ontology identified with GREAT using single closest gene. C-D) Heatmaps of enrichment in H3K27ac peaks in whole muscle and myonuclei in soleus and EDL, respectively.

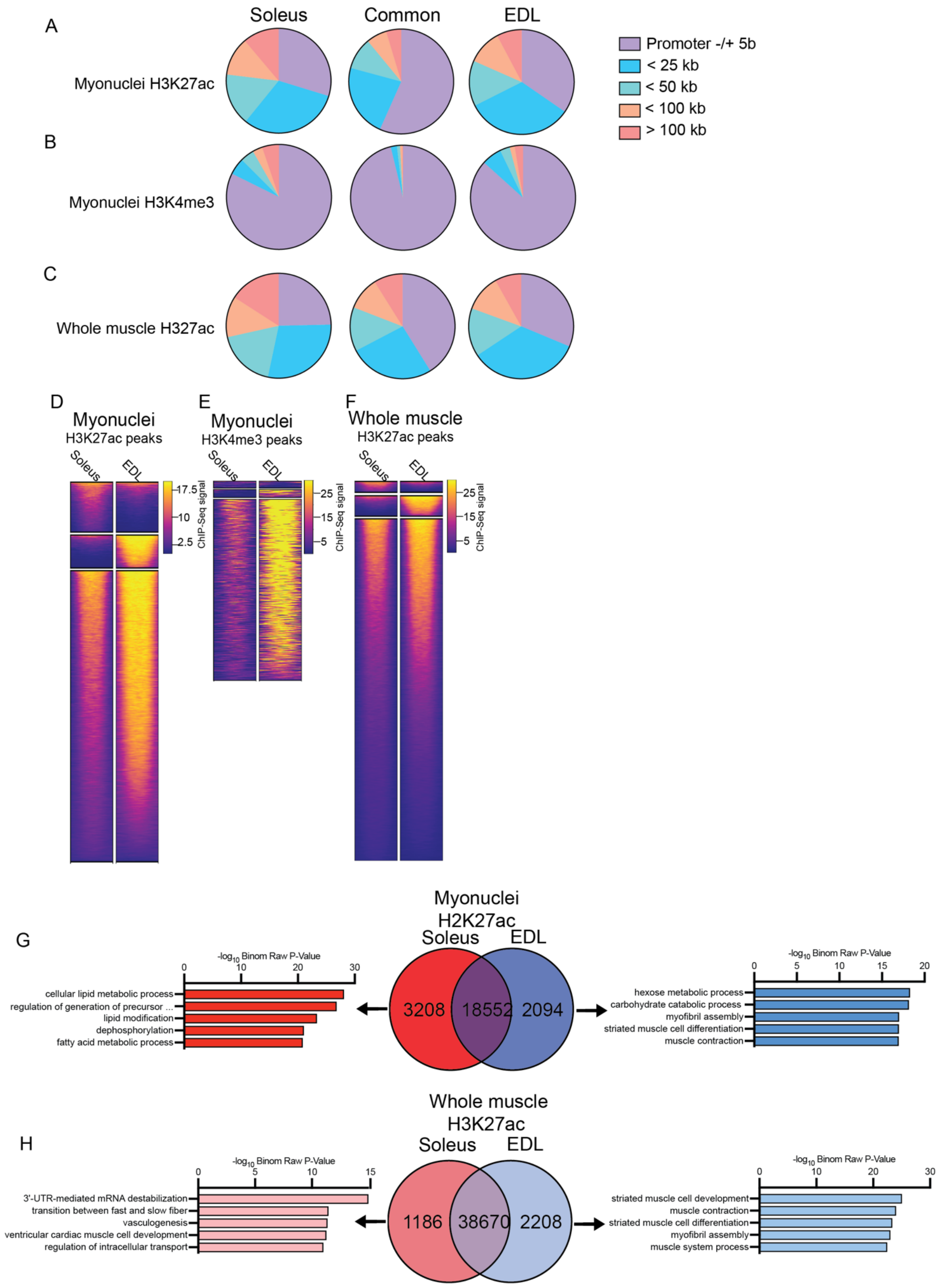


**Fig. S4: Epigenetic differences between whole muscle and myonuclei and muscle types.** A-C) Genomic distribution of differently enriched peaks between soleus and EDL. D-F) Heat map of the differently commonly enriched peaks for H3K27ac and H3K4me3 in myonuclei and H3K27ac in whole muscle. G-H) Venn diagram showing overlap between H3K27ac for soleus and EDL in myonuclei and whole muscle, respectively. Five most enriched gene ontologies for specific H3K27ac peaks are shown. For full list of ontologies see Table S2. Gene ontology identified with GREAT using single closest gene.

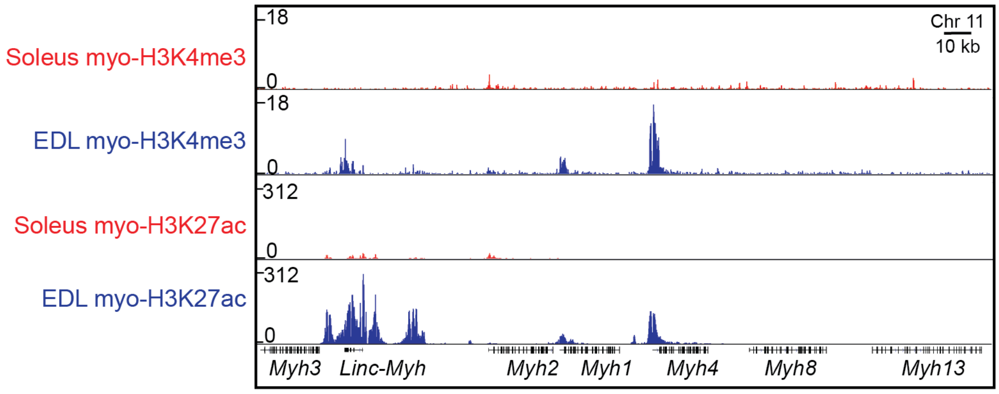


**Fig. S5: Only the mature skeletal muscle versions of myosin are enriched in adult muscle**. ChIP-Seq profiles of the myo-specific H3K4me3 and H3K27ac enrichment at the MyHC locus chromosome 11 encoding the embryonic myosin *Myh3* (MyHC-emb), the noncoding RNA *Linc-Myh*, the adult versions *Myh2* (MyHC-2A), *Myh1* (MyHC-2X), *Myh4* (MyHC-2B), neonatal myosin Myh8 (MyHC-neo) and the extraocular myosin *Myh13* (MyHC-EO).

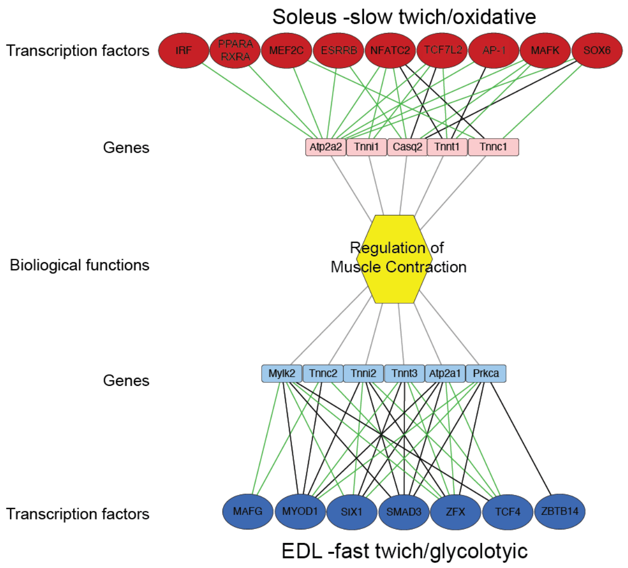


**Fig. S5. Soleus and EDL are enriched for different regulatory networks in muscle contraction.** Transcriptional regulatory network in muscle contraction for soleus (Top) and EDL (bottom). Closest genes with DE promoter for genes involved in regulation of muscle contraction inside 100 kb of the predicted binding motif. Color of the edges, black and green, indicates position of most significant motif prediction for factor at promotor region or distal regulatory enhancer region, respectively.
